# Angioplasty-induced epigenomic remodeling entails BRD4 and EZH2 hierarchical regulations

**DOI:** 10.1101/2020.03.12.989640

**Authors:** Mengxue Zhang, Bowen Wang, Go Urabe, Hatice Gulcin Ozer, Renzhi Han, K. Craig Kent, Lian-Wang Guo

## Abstract

Atherosclerosis is commonly treated with angioplasty which, however, evokes neointimal hyperplasia (IH) and recurrent stenotic diseases. Epigenomic investigation was lacking on post-angioplasty IH. The histone acetylation reader BRD4 and H3K27me3 writer EZH2 are potent epigenetic factors; their relationship is little understood. Through genome-wide survey in the rat angioplasty model, we studied BRD4 and EZH2 functional regulations involved in IH.

We performed chromatin immunoprecipitation sequencing (ChIPseq) using rat carotid arteries. While H3K27me3 ChIPseq signal prevalently intensified in balloon-injured (*vs* uninjured) arteries, BRD4 and H3K27ac became more enriched at *Ezh2*. Indeed, BRD4-siRNA or CRISPR-deletion of BRD4-associated enhancer abated the smooth muscle cell (SMC) expression of EZH2, and SMC-specific BRD4 knockout in *BRD4*^*fl/fl*^; *Myh11CreER*^*T2*^ mice reduced both H3K27me3 and IH in wire-injured femoral arteries. In accordance, post-angioplasty IH was exacerbated and mitigated, respectively, by lentiviral expression and pharmacological inhibition of EZH2/1; EZH2 (or EZH1) loss- and gain-of-function respectively attenuated and aggravated pro-IH SMC proliferative behaviors. Furthermore, while H3K27me3 ChIPseq signal magnified at *P57* and ebbed at *Ccnd1* and *Uhrf1* after injury, silencing either EZH2 or EZH1 in SMCs up-regulated *P57* and down-regulated *Ccnd1* and *Uhrf1*.

In summary, our results reveal an upsurge of EZH2/H3K27me3 after angioplasty, BRD4’s control over EZH2 expression, and non-redundant EZH2/1 functions. As such, this study unravels angioplasty-induced loci-specific H3K27me3/ac redistribution in the epigenomic landscape rationalizing BRD4/EZH2-governed pro-IH regulations.

## Introduction

Neointimal hyperplasia (IH) in the inner vascular wall obstructs blood flow engendering cardiovascular diseases. It not only occurs in atherosclerosis, but persists after recanalization of stenosed arteries, causing (re)stenosis. Drug-eluting stents/balloons are clinically deployed to impede post-angioplasty IH. However, they are unable to eradicate IH yet potentially thrombogenic, as disclosed by meta-analyses^1, 2^. The concerns culminated with three consecutive FDA warnings (in 2019) of increased mortality potentially associated with paclitaxel-eluting stents and balloons. A compelling agenda thus re-emerges to better understand IH pathogenesis for therapeutic improvement^3^.

Neointima is primarily formed by vascular smooth muscle cells (SMCs) that have transitioned to a migro-proliferative state^4^. With no DNA sequence changes, still the same genome – this SMC phenotypic/state transition is epigenetic in nature^5-7^. We previously reported that BRD4, a bromo/extraterminal family (BETs) protein, is a determinant of phenotypic transitions of SMCs and IH in rat models^6^. BRD4’s bromodomains read/bind histone acetyl-bookmarks while its C-terminus interacting with the transcription elongation complex. As such, BRD4 serves as a linchpin coupling lineage-related cis- and trans-factors to the transcription machinery, and localizes this assembly to specific genomic loci, thereby co-activating a select set of genes^8^. Inasmuch as BRD4 inhibition stymies SMC proliferation but not endothelial growth, as evidenced both in vitro and in vivo^3, 6, 9, 10^, targeting BRD4 appears to be an attractive anti-restenotic approach^3^. To this end, an imperative task is to understand BRD4-directed epigenetic mechanisms in the context of IH. While BRD4 is being intensively studied and clinically targeted in the cancer field^11^, SMC-specific BRD4 epigenomic/epigenetic regulations in IH pathogenesis remain poorly explored^12^.

In contrast to BRD4’s function of co-activating transcription^8^, enhancer of zeste homologs (EZH2 and EZH1 isoforms) catalyze methylation at H3K27 leading to transcriptional repression^13, 14^. EZH2 recently emerged as a driver of cell state transitions such as cancer cell proliferation and migration, yet reports on EZH1 are limited and discrepant regarding its redundancy with EZH2^13, 15^. Pharmacological evidence from our^16^ and other groups^17, 18^ supports an IH-mitigating effect of pan-EZH1/2 inhibition, underscoring the importance of dissecting their functional roles and the mechanisms governing their expression levels. Moreover, BRD4 as an acetylation reader and EZH2 as a methylation writer are seemingly unrelated; their relationship has been overlooked^19^, especially in vascular pathogenesis. However, it is important to explore a functional connection between BRD4 and EZH2, both cell identity/state determinants^20^ and targets of active clinical trials^14^.

Epigenomic studies pertaining to IH have been limited and mostly confined to cell cultures^7, 21, 22^, which as oversimplified systems inevitably produce inaccurate information for interpreting in vivo pathogenic mechanisms^12, 23^. Here we performed ChIPseq directly using rat carotid arteries that underwent angioplasty injury-induced IH. We observed a prominent injured-vs-uninjured genome-wide upsurge of H3K27me3, a repression mark^20^. This was, at first sight, counter-intuitive since conventional views regard transcriptional activation as the prevailing event following injury in the IH model. Yet, rationalizing this result, angioplasty promoted BRD4/H3K37ac genomic re-localization to *Ezh2*. Indeed, BRD4 silencing and BRD4-associated enhancer deletion indicated that BRD4 governed EZH2 expression in SMCs, which in turn repressed cytokinetic inhibitors such as P57. As such, this study uncovered the functional interplay between acetylation reader BRD4 and H3K27me3 writer EZH2 closely involved in IH pathogenesis.

## Results

### Angioplasty in rat carotid arteries induces changes in ChIPseq peaks

There has been a paucity of knowledge on in vivo genome-wide epigenetic regulations during angioplasty injury-induced neointima formation. An authentic model to mimic clinical angioplasty, namely, balloon angioplasty of rat common carotid arteries^3, 6^, confers an opportunity for ChIPseq studies using artery tissues that undergo IH. It could be more convenient to do ChIPseq using cultured cells, however, important information of in vivo pathological processes would be inevitably missed. We therefore performed ChIPseq using balloon-injured (and uninjured contralateral) rat carotid arteries. Typically, following balloon angioplasty, the endothelial inner lining of the artery is removed, and SMCs – the major constituent cell type in the artery wall – become exposed to various stimuli such as platelet derived growth factor (PDGF), and undergo migro-proliferative phenotypic transitions forming neointima. We collected the arteries at post-angioplasty day 7 as this is the peak time of a myriad of pro-IH molecular and cellular events^24, 25^. Epigenomic marks involved in transcriptional activation, including BRD4, H3K27ac, and H3K4me1^26^, were chosen for ChIP experiments. We also included H3K27me3, a mark of transcriptional repression^27^. The distribution of ChIPseq peaks around the TSSs of over 24,000 genes is shown in Figure 1A. Hierarchical clustering of the peaks shows that majority of the BRD4, H3K27Ac and H3K4me1 peaks co-localize around TSS regions, and this is mutually exclusive with H3K37me3 peaks. Distribution of individual ChIPseq peak scores is presented as bean plots in Figure 1B. H3K27ac and H3K27me3 signal shows augmentation after injury. When plotted to show the peaks with increased or decreased intensity using a 2-fold change as cutoff (Figure 1C), there appears a prevailing shift of H3K27me3 ChIPseq peaks to the side of boosted intensity. Injury-induced ChIPseq signal augmentation also occurred to H3K27ac. These results reveal remarkable angioplasty-induced H3K27me3/ac genome-wide remodeling in vivo in rat carotid arteries.

**Figure 1.**
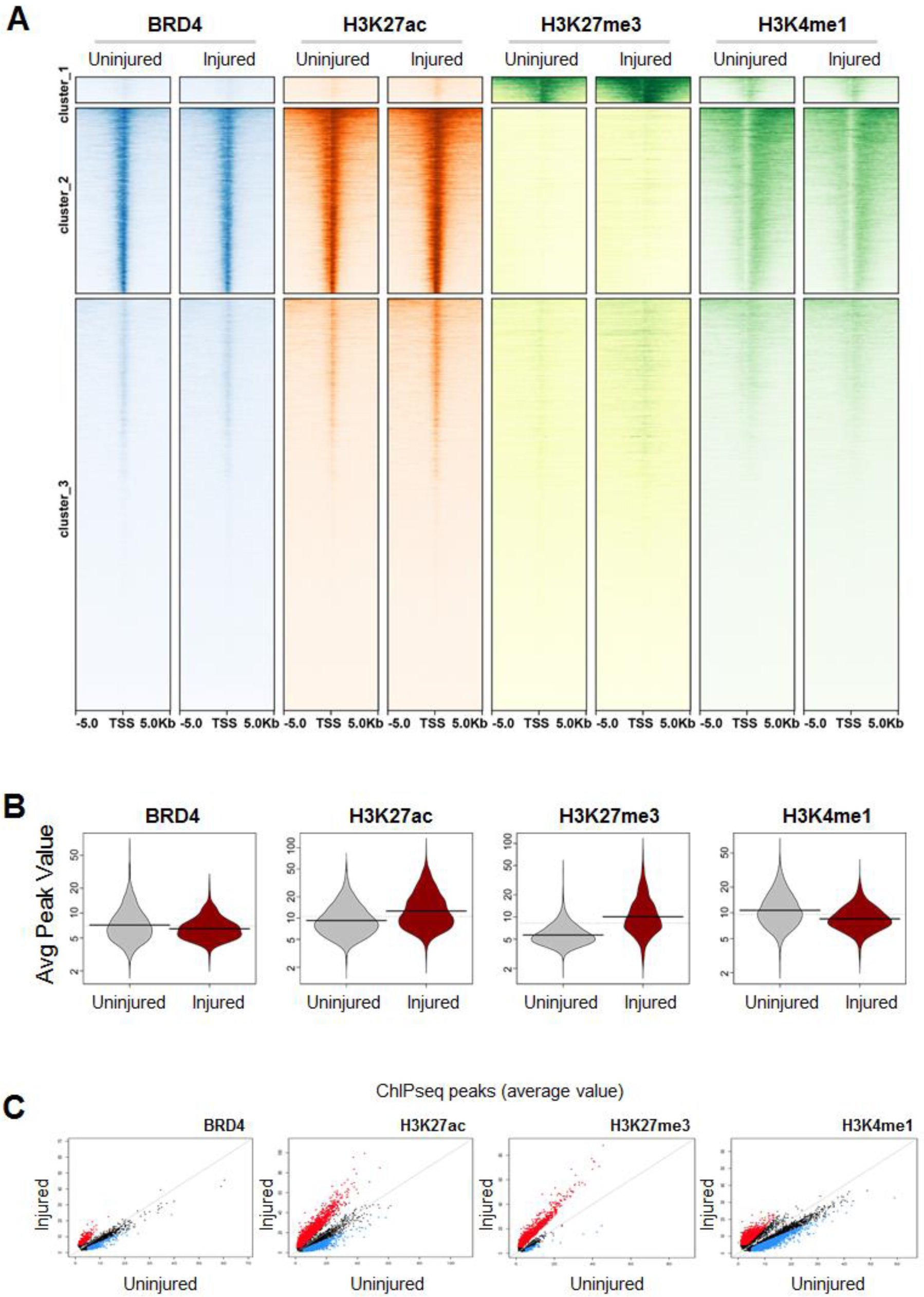
Injury-induced changes of ChIPseq peaks in rat carotid artery tissues. Balloon-injured rat left common carotid arteries and contralateral arteries (uninjured control) were collected at day 7 post angioplasty and snap frozen until use for ChIPseq. **A**. Heatmaps of ChIPseq peak density for BRD4, H3K27ac, H3K27me3, and H3K4me1. ChIPseq signal anchors 10 kb center region with 5 kb flanking on either side of the transcription start site (TSS) of over 24000 genes. Three clusters show the main pattern of co-localization of the ChIP-seq signal and non-overlap between H3K27ac and H3K27me3. Note increased (injured-*vs*-uninjured) H3K27me3 ChIPseq signal mainly in Cluster-1. **B**. Distribution of transcript abundance of the genes associated with BRD4 and the three histone marks. **C**. ChIPseq intensity changes in injured (*vs* uninjured) arteries. Red, increased intensity; blue, decreased intensity; a cutoff of 2-fold change of read intensity was used. Note the prevailing H3K27me3 ChIPseq signal increase after injury.

### BRD4 and H3K27ac enrich at the EZH2 gene in injured arteries; BRD4 governs EZH2 expression in SMCs

Since EZH2 is the primary methyltransferase that deposits H3K27me3^27^, the striking H3K27me3 upsurge due to angioplasty led us to investigate the underlying epigenetic regulations. As observed in our data^16^ and recently reproduced by others^17, 18^, pharmacological evidence implicated EZH2 as potentially important in neointima formation. Interestingly, both BRD4- and H3K27ac-associated ChIPseq peaks enriched at *Ezh2*, more in injured *vs* uninjured arteries (Figure 2A). Angioplasty also induced BRD4/H3K27ac enrichment at *Nrp2* (Figure 2B), a recently identified pro-IH gene markedly up-regulated by angioplasty in the same rat IH model^28^. As such, *Nrp2* provided an ideal positive control validating our methodology and ChIPseq results. In our recent report, BRD4 was identified as an epigenetic determinant of IH which was highly upregulated at post-angioplasty day 7. We were thus intrigued as to whether BRD4, a histone acetylation reader, regulates the expression of EZH2 which is a histone methylation writer. Through siRNA-mediated gene silencing in SMCs, we found that BRD4, but not other BETs (BRD2 or BRD3), controlled EZH2 mRNA and protein levels (Figure 2, C and D). Since BRD4 is an epigenomic mark of transcription enhancers^26^, we next explored the function of BRD4-associated enhancers for EZH2 expression. As shown in Figure 2 (E and F), CRISPR-mediated genomic deletion of an enhancer region reduced EZH2 mRNA and protein. These results indicate that BRD4 governs the expression of *Ezh2* in SMCs, an action likely involving its association with enhancers.

**Figure 2.**
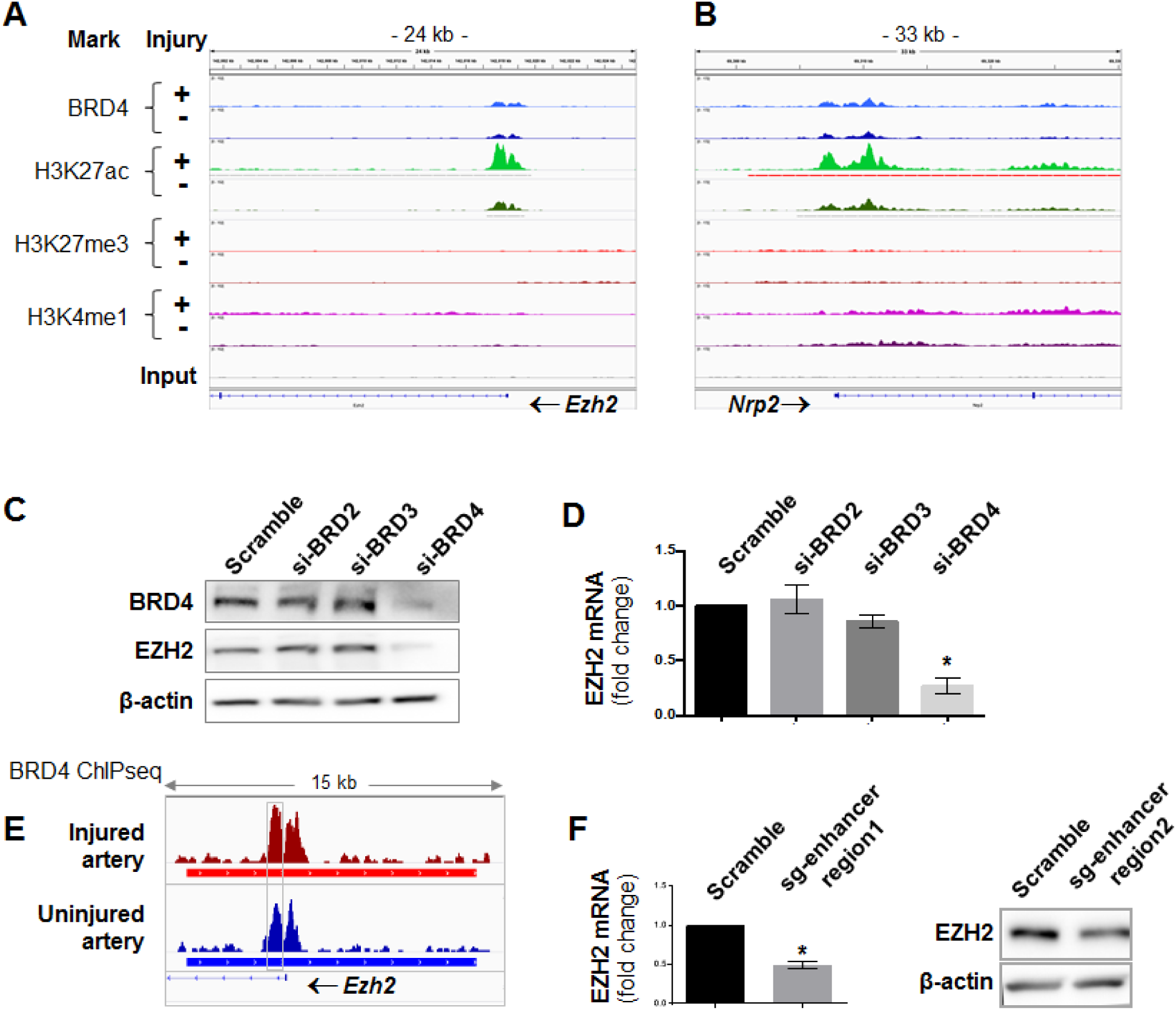
Increase of injured-vs-uninjured BRD4/H3K27ac enrichment at Ezh2. **A** and **B**. Comparison of ChIPseq binding density (near *Ezh2*) between injured (+, light color) and uninjured (-, dark color) arteries. The profiles for *Nrp2*, which is known as upregulated in balloon-injured rat carotid arteries^28^, are presented for positive control to validate the ChIP methodology and data. Non-specific input indicates very low background noise. **C** and **D**. Effect of BRD4 silencing on EZH2 expression. BRD2, BRD3, or BRD4 was silenced with their specific siRNAs (validated in our recent reports)^6, 10^. Rat aortic SMCs were cultured, starved for 6h, then transduced with BRD2,3,4 siRNA overnight, recovered for 24h and 48h before RNA and protein extraction.EZH2 protein and mRNA were measured with Western blot and qRT-PCR (normalized by ΔΔCT-log2) assays. Quantification: Mean ± SEM; n =3 independent experiments; one-way ANOVA with Bonferroni test, *P<0.05 compared to the scrambled-sequence siRNA control. **E**. BRD4 ChIPseq binding density focusing on *Ezh2*. Red and blue bars mark enhancers. Box highlights an enhancer region where ChIPseq intensity increased in injured vs uninjured arteries. **F**. Effect of CRISPR-mediated enhancer region deletion on EZH2 expression. sg, short guide RNA. Quantification: Mean ± SEM; n =3 independent experiments; one-way ANOVA with Bonferroni test, *P<0.05.

### SMC lineage-specific BRD4 deletion in mice reduces IH and H3K27me3 in wire-injured femoral arteries

To examine the in vivo function of the BRD4/EZH2 regulatory axis, we first performed conditional knockout of BRD4 and IH-inducing wire injury (Figure 3, A and B). Mice were cross-bred with the strains of *Brd4*^*fl/fl*^ and *Myh11-CreER*^*T2*^. Tomaxifen-containing chow was fed to *Brd4*^*fl/fl*^; *Myh11-CreER*^*T2*^ mice to induce SMC-specific BRD4 knockout (KO) followed by wire injury and collection of femoral arteries for histology (Figure 3B). As shown in Figure 3 (C and D), IH (normalized as I/M ratio) was drastically reduced in homozygous BRD4 KO mice, either compared to the wild type (*Brd4*^*+/+*^) or heterozygous KO (*Brd4*^*+/-*^) animals. This result concurs with our previous reports using pharmacological and shRNA approaches^6, 10^. Thus, the IH data is significant not only because it validates the SMC-specific KO experimental setting, but it represents the first-time demonstration of the SMC-specific role for BRD4 in neointimal development. We then measured H3K27me3 on artery cross sections as a surrogate of EZH2’s writer function (Figure 3, E and F). H3K27me3 was markedly lower in *Brd4*^*fl/fl*^; *Myh11-CreER*^*T2*^ *vs Brd4*^*fl/fl*^ mice, as observed in the medial and neointimal layers where the major constituent cells are SMCs. This and other results (including Figures 1 and 2) together suggest a BRD4 > EZH2 > H3K27me3 hierarchical regulation, especially considering that H3K27me3 is known as as deposited primarily by EZH2^27^.

**Figure 3.**
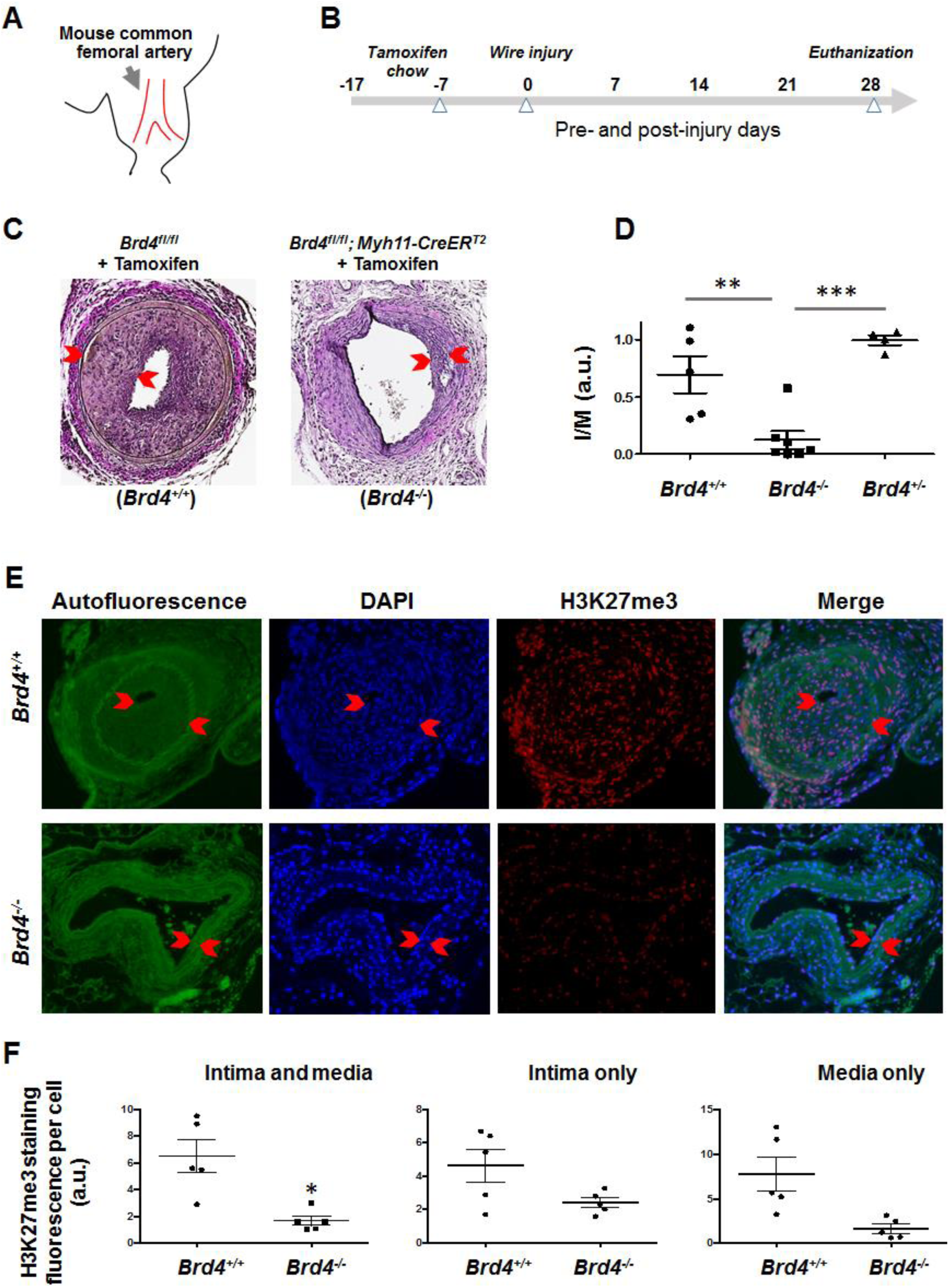
SMC-specific BRD4 knockout reduces IH in wire-injured mouse femoral arteries. **A**. Cartoon showing mouse common femoral artery where wire injury was made to induce IH. **B**. Diagram indicating the time line for tamoxifen feeding, wire injury, and tissue collection. **C** and **D**. Comparison of IH between WT (*Brd4*^*fl/fl*^) and induced SMC-specific BRD4 knockout (*Brd4*^*fl/fl*^; *Myh11-CreER*^*T2*^) mice. Neointima thickness is indicated between arrow heads. IH is normalized as intima/media area ratio. **E** and **F**. Comparison of H3K27me3 (immunostaining) between WT and BRD4 knockout mice. Neointima is indicated between arrow heads. Immunofluoresence was normalized with cell number. Quantification (in D and F): Mean ± SEM; n (number of mice) is indicated by the data points in scatter plots; one-way ANOVA with Bonferroni test, *P<0.05, **P<0.01, ***P<0.001; a.u. arbitrary unit.

### EZH2 and EZH1 each promotes IH in balloon-injured rat carotid arteries

In addition to EZH2, the EZH1 isoform also methylates H3K27^13^. However, EZH1 is much less studied. Its specific role in IH was not known, since previous pharmacological studies using a pan inhibitor of EZH1/2 failed to distinguish their individual contributions to IH^17, 18^. To address this knowledge gap, we first observed that pre-treatment of SMCs with JQ1, a bromodomain blocker binding BRD4 (and other BETs)^11^ abrogated PDGF-stimulated mRNA expression of not only EZH2 but also EZH1 (Figure 4A). We then determined their expression after balloon angioplasty in rat carotid arteries. The data showed a continuous increase of EZH2 throughout the 14-day time course (Figure 4B). EZH1 was upregulated as well in the later stage (day 14) albeit with an initial dip observed for day 3. To delineate the specific roles of the EZH isoforms in neointimal development, we performed gain- and loss-of-function experiments in vivo using the rat carotid artery angioplasty model. We found that compared to the GFP control, increasing either EZH1 or EZH2 via lentiviral gene transfer to the injured artery wall (Figure 4C) exacerbated IH and restenosis (lumen narrowing) (Figure 4, D and E). Consistently, perivascular local treatment of injured arteries with the pan-EZH1/2 inhibitor UNC1999 (Figure 4F) diminished IH and enlarged the lumen (Figure 4, G and H). While clarifying the pre-conceived role of EZH2^17, 18^, our results indicate that EZH1 also plays a positive role in neointima development, as demonstrated in the rat angioplasty model.

**Figure 4.**
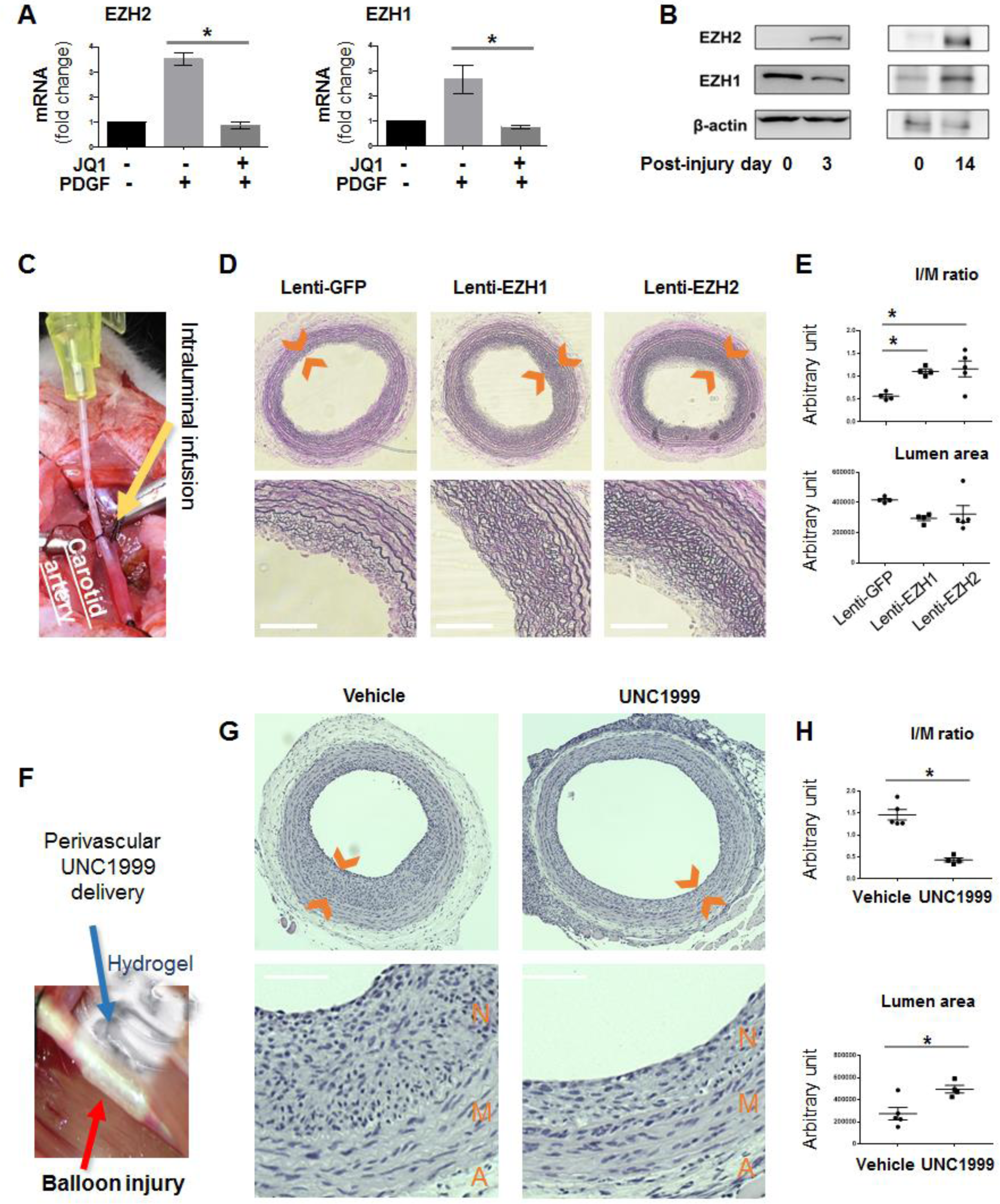
Effect of EZH1 or EZH2 gain- or loss-of-function on IH in balloon-injured rat carotid arteries. **A**. Effect of blocking BRD4 function (with JQ1) on the expression of EZH2 and EZH1. Rat primary aortic SMCs were cultured, starved, pre-treated with vehicle (DMSO) or the pan-BET inhibitor JQ1 (1 µM) for 2h, and then added with PDGF-BB (final 20 ng/ml) and cultured for another 1h prior to cell harvest for qRT-PCR. Mean ± SEM; n =3 independent experiments; one-way ANOVA with Bonferroni test, *P<0.05. **B**. Up-regulation of EZH2 and EZH1 in balloon-injured rat carotid arteries. Arteries were collected at post-injury day 3 and day 14 for Western blot analysis. Day 0 refers to uninjured control. **C**. Picture illustrating intraluminal infusion of lentivirus to express a gene in the balloon-injured artery wall. A cannula (yellow device) connected to a syringe was used to inject lentivirus to the injured carotid artery lumen for infusion into the denuded artery wall. **D** and **E**. Gain of function. Overexpression of GFP (control), EZH1, or EZH2 in the injured artery wall was accomplished via intraluminal infusion of lentivirus. Arteries were collected at post-injury day 14 for histology. Neointima is shown between arrow heads (D). Quantification (E): Mean ± SEM; n (number of rats) is indicated by data points in scatter plots; one-way ANOVA with Bonferroni test, *P<0.05. **F**. Picture depicting perivascular drug delivery. The inhibitor drug (UNC1999) dispersed in hydrogel (AK12, liquid on ice and paste-like at body temperature) can be applied around an injured carotid artery (after pulling out the balloon). **G** and **H**. Loss of function. The pan-EZH1/2 inhibitor UNC1999 was administered as explained in F. Arteries were collected at post-injury day 14. Quantification: Mean ± SEM; n (number of rats) is indicated by data points in scatter plots; one-way ANOVA with Bonferroni test, *P<0.05.

### EZH2 and EZH1 are functionally non-redundant in promoting the migro-proliferative SMC phenotypic transition

To further dissect the EZH2/1-mediated functional mechanisms, we used the PDGF-induced cellular model that exhibits salient pro-IH migro-proliferative SMC phenotypes^6^. Pre-treatment with the pan-EZH1/2 inhibitor UNC1999 concentration-dependently inhibited PDGF-induced SMC proliferation (Figure 5A) and migration (Figure 5B). Furthermore, in an isoform-specific manner, silencing either EZH2 or EZH1 with shRNA markedly inhibited PDGF-induced SMC proliferation and migration (Figure 5, C-E). This result for the first time revealed that EZH2 and EZH1 were non-redundant in promoting the pro-IH SMC behaviors. This is interesting given that redundancy of EZH2 and EZH1 was commonly reported in other biological contexts^15, 27^. In further support of this conclusion, lentivirus-mediated gain-of-function experiments indicated that increasing either EZH2 or EZH1 aggravated SMC proliferation and migration (Figure 5, F-H). Taken together, the in vitro (Figures 5 and 6) and in vivo (Figure 4) results indicate that EZH2 and EZH1 are functionally non-redundant in PDGF-induced SMC phenotypic transitions, each playing a positive role in neointimal development.

**Figure 5.**
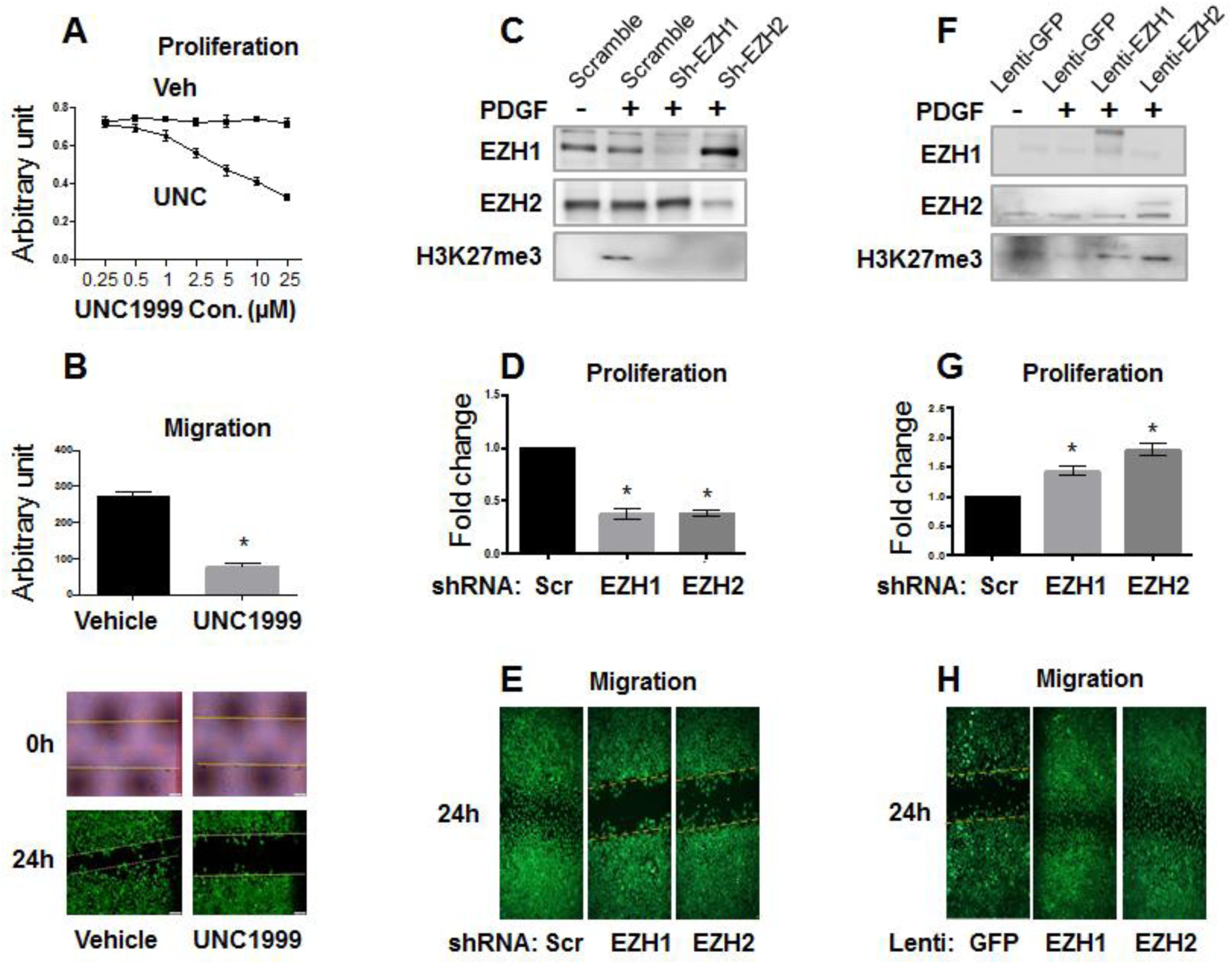
Effect of EZH1 or EZH2 gain- or loss-of-function on SMC proliferation/migration. MOVAS cells were cultured, starved, pre-treated with vehicle (DMSO) or the pan-EZH1/2 inhibitor UNC1999 (5 µM) for 2h, and then stimulated with PDGF-BB (final 20 ng/ml). For lentivirus-mediated overexpression or silencing, cells were transduced with lentivirus overnight, recovered for 24h, and then starved for 6h prior to adding PDGF-BB. After 24h, 72h, and 96h of stimulation with PDGF-BB, cells were harvested for proliferation, qRT-PCR, and Western blot assay, respectively. Quantification: Mean ± SEM; n =3 independent experiments; one-way ANOVA with Bonferroni test, *P<0.05. **A**. Effect of UNC1999 pretreatment on SMC proliferation (Cell TiterGlo assay). Cells were harvested after 24h stimulation with PDGF-BB. **B**. Migration (scratch assay). To measure migration, cells were pictured at the beginning (0h) and end (24h) of PDGF-BB stimulation. Statistics: student t-test, *P<0.05. **C-E**. Loss of function. Silencing efficiency is indicated by Western blots (C). For proliferation (D) and migration (E) assays, cells were harvested at 72h or pictured at 24h after PDGF-BB stimulation, respectively. Scr, scrambled. **F-H**. Gain of function. EZH1 or EZH2 overexpression is shown in Western blots (F). For proliferation (D) and migration (E) assays, cells were harvested at 72h or pictured at 24h after PDGF-BB stimulation, respectively.

**Figure 6.**
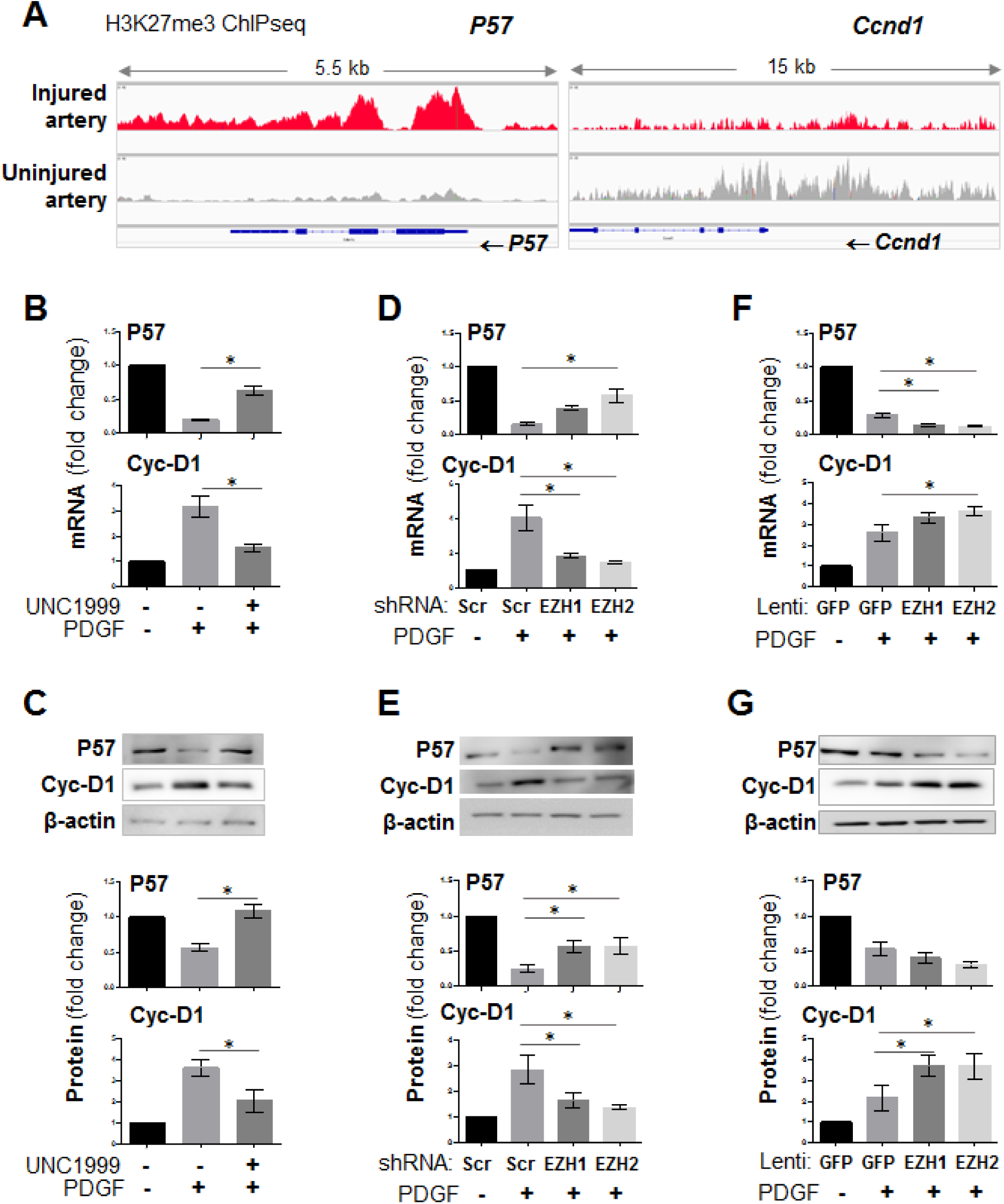
H3K27me3 ChIPseq binding density at P57 and cyclin-D1 and their regulation by EZH1 and EZH2. MOVAS cells were cultured, starved, pre-treated with vehicle (DMSO) or the pan-EZH1/2 inhibitor UNC1999 (5 µM) for 2h, and then stimulated with PDGF-BB (final 20 ng/ml). For lentivirus-mediated overexpression or silencing, cells were transduced with lentivirus overnight, recovered for 24h, and then starved for 6h prior to adding PDGF-BB. Cells were harvested 24h or 48h after stimulation with PDGF-BB, for qRT-PCR and Western blot assay, respectively. Quantification: Mean ± SEM; n =3 independent experiments; one-way ANOVA with Bonferroni test, *P<0.05. **A**. H3K27me3 ChIPseq binding density at *Cdkn1c* (P57) and *Ccnd1* (cyclin-D1) in balloon-injured (red) and uninjured (gray) artery tissues. **B** and **C**. Effect of pan-EZH1/2 inhibition on P57 and cyclin-D1 expression. **D** and **E**. Effect of EZH1 or EZH2 silencing on P57 and cyclin-D1 expression. **F** and **G**. Effect of EZH1 or EZH2 overexpression on P57 and cyclin-D1 expression.

### Angioplasty induces H3K27me3 redistribution from Ccnd1 to P57; both are regulated by EZH2 and EZH1 in SMCs

H3K27me3 is generally known to be transcriptionally repressive, providing us clue to track down target genes of the EZH2/1 writer function known as transcription repression^15^. We thereby revisited the ChIPseq data for H3K27me3. In stark contrast to uninjured control, H3K27me3 in injured arteries highly enriched at *P57* (a.k.a. *Cdkn1c*), a bona fide inhibitor of cell proliferation/migration^24^. The opposite occurred to *Ccnd1* (cyclin D1), a potent pro-proliferative/migratory cytokinetic factor (Figure 6A). Validating the ChIPseq data, the pre-treatment of SMCs with UNC1999 restored *P57* expression yet inhibited *Ccnd1* expression, at both mRNA and protein levels in the presence of PDGF-BB (Figure 6, B and C). To dissect the individual EZH2 or EZH1 function, we applied lentiviral expression of shRNA or transgene to specifically decrease or increase EZH2 or EZH1. The mRNA and protein data together demonstrated that either EZH1 or EZH2 loss-of-function partially reinstated *P57* expression that was hampered by PDGF, while blocking the PDGF induction of *Ccnd1* expression (Figure 6, D and E). In accordance, the gain-of-function experiments led to opposite results (Figure 6, F and G). Therefore, the collective results reiterate the non-redundancy of the two EZH isoforms in the context of pro-IH SMC pathobiology.

### Angioplasty induces H3K27me3 substitution by H3K27ac at Uhrf1, which is found here as a novel target of EZH1/2 transcriptional regulation

Up to this point, existing literature evidence for EZH2 regulating *P57*^14^ and our consistent new results of epigenomic survey and molecular elaboration had conferred a reliable “compass” to navigate the epigenomic landscape remodeled due to angioplasty. It was on this basis that we further quested for novel target genes that responded to H3K27me3 dynamics. As presented above (Figure 1C), angioplasty induced prevailing rise of H3K27m3 ChIPseq peak intensity for the majority of genes, in a conspicuous contrast to diminution of the H3K27m3 signal for only a small number of genes (Figure 1C) among which we found *Uhrf1* as top-ranked in addition *to Ccnd1*. Further attracting our interest, ChIPseq signal for H3K27ac at *Uhrf1* magnified after injury (Figure 7A). Consistently, recent reports indicated that UHRF1 functionally associates with histone methylation and acetylation^29^, and its upregulation in injured arteries promotes SMC proliferation and IH^30^. Interestingly, *Ezh2* and *Uhrf1* were reported to be in the same gene network both promoting keratinocyte self-renewal^31^, yet their epigenomic relationship was not known. Here we found that while EZH2 (or EZH1) loss-of-function reduced, their gain-of-function increased *Uhrf1* mRNA (Figure 7B). Thus, we identified UHRF1 as a novel target of EZH1/2 functional regulation, a finding consistent with the recently reported positive role of UHRF1 in SMC proliferation/migration and injury-induced IH^30^.

**Figure 7.**
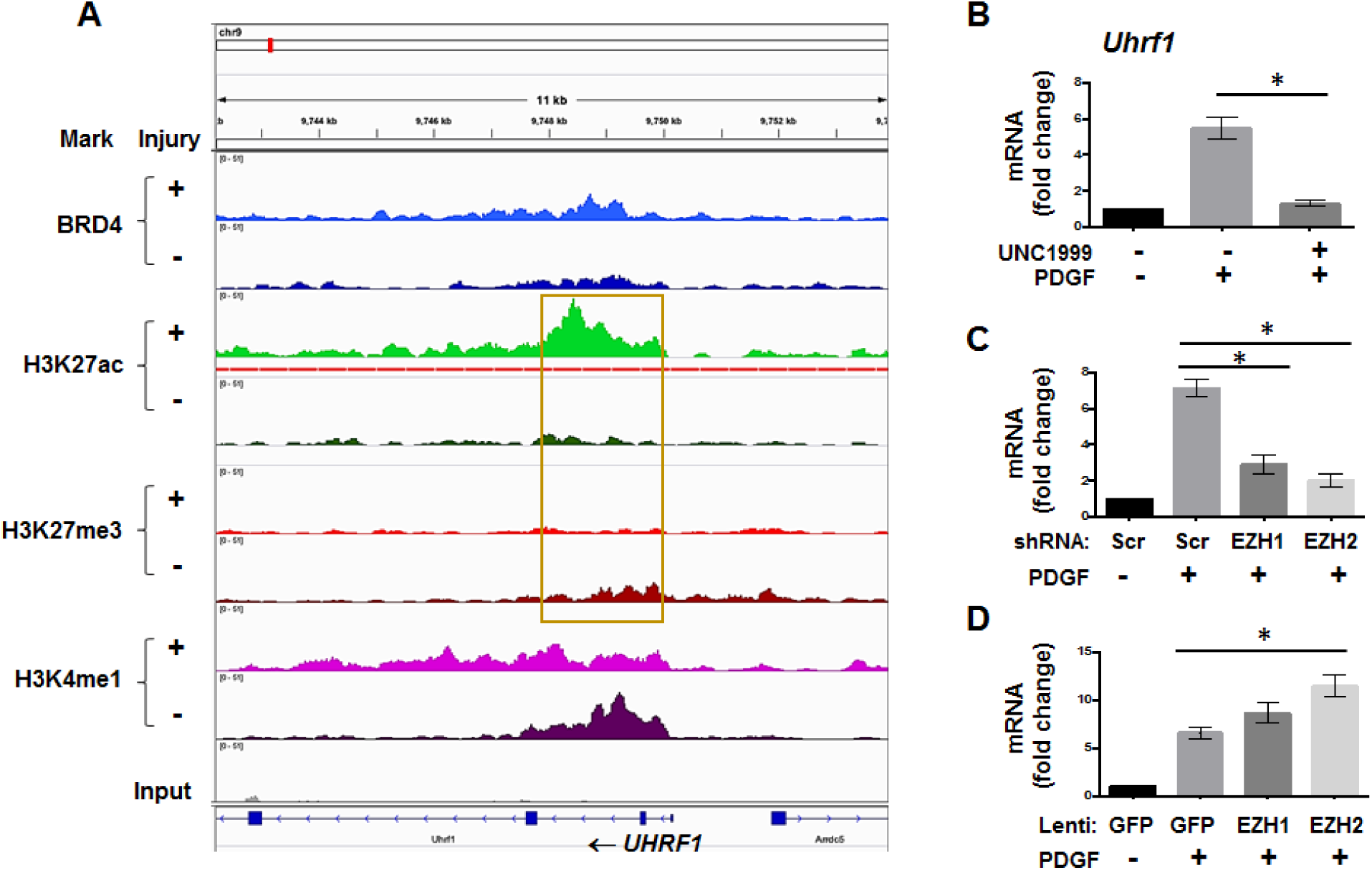
H3K27me3 ChIPseq binding density at Uhrf1 and its regulation by EZH1 and EZH2. MOVAS cells were cultured, starved, pre-treated with UNC1999 or transduced with lentivirus, stimulated with PDGF-BB, and assayed, as described for Figure 6. Quantification: Mean ± SEM; n =3 independent experiments; one-way ANOVA with Bonferroni test, *P<0.05. **A**. H3K27me3 ChIPseq binding density at *Uhrf1* in balloon-injured (red) and uninjured (gray) artery tissues. **B**. Effect of pan-EZH1/2 inhibition on UHRF1 expression. **C**. Effect of EZH1 or EZH2 silencing on UHRF1 expression. **D**. Effect of EZH1 or EZH2 overexpression on UHRF1 expression.

## Discussion

Epigenetic remodeling is increasingly recognized as crucial in cardiovascular pathologies such as IH^12^. The histone acetylation reader BRD4 and H3K27me3 writer EZH2 are powerful epigenetic regulators, as reported for proliferative diseases^11, 14, 26^ and implicated for IH. While rapidly growing, epigenetic studies concerning IH are mostly reliant on cell cultures^21, 22^. Here we report in vivo genome-wide epigenetic survey in arteries that undergo injury-induced IH. We found surging rather than fading H3K27me3 ChIPseq peak intensity after IH-inducing angioplasty. In accordance, BRD4 enrichment at *Ezh2* increased after angioplasty, and BRD4 governed EZH2 expression in SMCs. Furthermore, not only EZH2 but also EZH1 promoted IH and proliferation/migration of SMCs, each associated with repression of P57 and de-repression of cyclin-D1 expression. Thus, the combined in vitro and in vivo results revealed previously unknown BRD4/EZH2 regulations involved in IH-promoting epigenetic remodeling.

We previously reported that BRD4, a transcription co-activator^11^, dramatically increased in the neointima and strongly promoted IH in either an angioplasty^6^- or vein grafting-induced rat model^10^. However, we did not realize an inner connection between BRD4 and EZH2 when we first observed an IH-mitigating effect of a pan-EZH inhibitor^16^. It then stuck us as a surprise that arterial injury induced upsurge of ChIPseq peaks for H3K27me3 (Figure 1C), a well-established transcriptional repression mark^27^. The traditional view has it that activation rather than repression of numerous genes and pathways is prevailing after angioplasty^24^, which injures the artery and denudes endothelium thereby exposing medial SMCs to a myriad of stimuli that trigger SMC proliferative state transitions and IH^3^. On this knowledge basis, it was counterintuitive to see that the majority of genes were associated with rising H3K27me3 ChIPseq peaks (injured *vs* uninjured) whereas only a small number of genes were on the opposite side (Figure 1C).

While tracing the factor(s) underlying enhanced H3K27me3, which is known to be deposited primarily by EZH2^13^, we noticed greater BRD4 and H3K27ac enrichment at *Ezh2* in injured *vs* uninjured arteries. This data led us to investigate whether BRD4 regulated EZH2 expression given that both BRD4 and H3K27ac are enhancer marks typically associated with transcriptional activation^26^. Indeed, the gene silencing data unequivocally indicated that BRD4, but not the other two BETs (BRD2 and BRD3) was the determinant of EZH2 expression. Moreover, enhancer deletion reduced EZH2 as well. These results fit the current working model that BRD4 rallies multiple factors (including enhancers) to facilitate transcription elongation by reading/binding H3K27ac while coupling with the transcription machinery^8, 11^. Because BRD4 and EZH2 each potently promotes IH, as demonstrated by our conditional knockout and gene transfer in vivo experiments presented herein, this BRD4/EZH2 axis is mechanistically important. In support of our finding, enhanced efficacy of combined BETs and EZH2 inhibitors has been reported in cancer research^32^. Moreover, a recent report showed that BRD4 regulated EZH2 expression by indirectly upregulating c-myc in cancer cells^19^, although it was not addressed as to whether epigenetic mechanisms were involved.

Now that BRD4 was identified as an upstream epigenetic determinant of EZH2 expression, we further elaborated EZH2 downstream functions. Interestingly, our data indicated that EZH2 and EZH1 each played an important role in promoting SMC proliferation/migration and IH. This non-redundancy was somewhat unexpected. As a much less studied isoform, EZH1’s function in IH was not known. Whereas EZH1 was deemed redundant to EZH2 in various tissues, e.g. tumor and skin^15, 27^, their non-redundant roles were recently found in development^13^. If EZH1/2 redundancy had occurred in our study, silencing one would have been compensated for by the unsilenced other isoform. Apparently, our data indicated that it was not the case. This finding could be instructive for therapeutic purposes, e.g. an inhibitor selective to either of the EZH isoforms could be effective in mitigating IH, thus conferring options in case inhibiting one isoform would cause unwanted complications such as impaired cardiac development^13^. This proposition requires future experiments to test as highly isoform-selective inhibitor drugs are currently unavailable^14^.

In pursuit of regulatory mechanisms downstream of EZH1/2, our ChIPseq data provided an important clue; that is, H3K27me3 heavily enriched at the P57 gene in injured arteries *vs* uninjured control. Indeed, loss- and gain-of-function experiments with SMCs confirmed that EZH1 and EZH2 repressed the expression of P57, a bona fide cell cycle inhibitor against SMC migro-proliferative behaviors^24^. While P57 is exemplary, other cell cycle inhibitors such as P27 followed suit. These turned out to be only part of the story, as the factors antagonizing P57, represented by cyclin-D1, were elevated due to EZH1/2 gain-of-function. More interestingly, UHRF1 belonged to this group. Other than a cytokinetic factor, UHRF1 is a multifunctional epigenetic reader without a previously recognized functional connection with EZH2^29^. Very recently, UHRF1 was reported to play a critical role in IH^30^, and in another study found to associate with both methylation and acetylation histone marks^29^. Thus, its potential functional interplay with BRD4 and EZH2 in the IH context deserves a new project to explore.

At this point, the seemingly paradoxical post-angioplasty upsurge of repressive ChIPseq signal (i.e. H3K27me3) could be rationalized (refer to the schematic in Figure 8). As arterial injury resets the epigenome, heightened BRD4/H3K27ac enrichment at *Ezh2* upregulates EZH2 and its enzymatic product H3K27me3, which represses the expression of cell cycle inhibitors. On the other hand, reduced H3K27me3 at cell cycle activators de-represses their expression. In either case, SMC proliferation and IH are exacerbated. Likely through this “Yin-Yang” regulation, BRD4 and EZH2 together effect a “double whammy” to efficiently propel pro-IH SMC proliferation. In a full view of the epigenomic landscape, this action appears to be executed at least in two modes. In a lateral mode, H3K27me3 relocates, leading to its gain and loss at different loci, e.g. more at *P57* and less at *Ccnd1*. In an on-site fashion, H3K27ac substitutes H3K27me3 at the same gene, as exemplified by *Uhrf1*. H3K27ac stakes off H3K27me3 and loss of H3K27me3 vacates the site for H3K27ac^20^, either benefiting *Uhrf1* transcription. As such, the reconstructed epigenomic landscape may have paved a way for accelerated IH.

**Figure 8.**
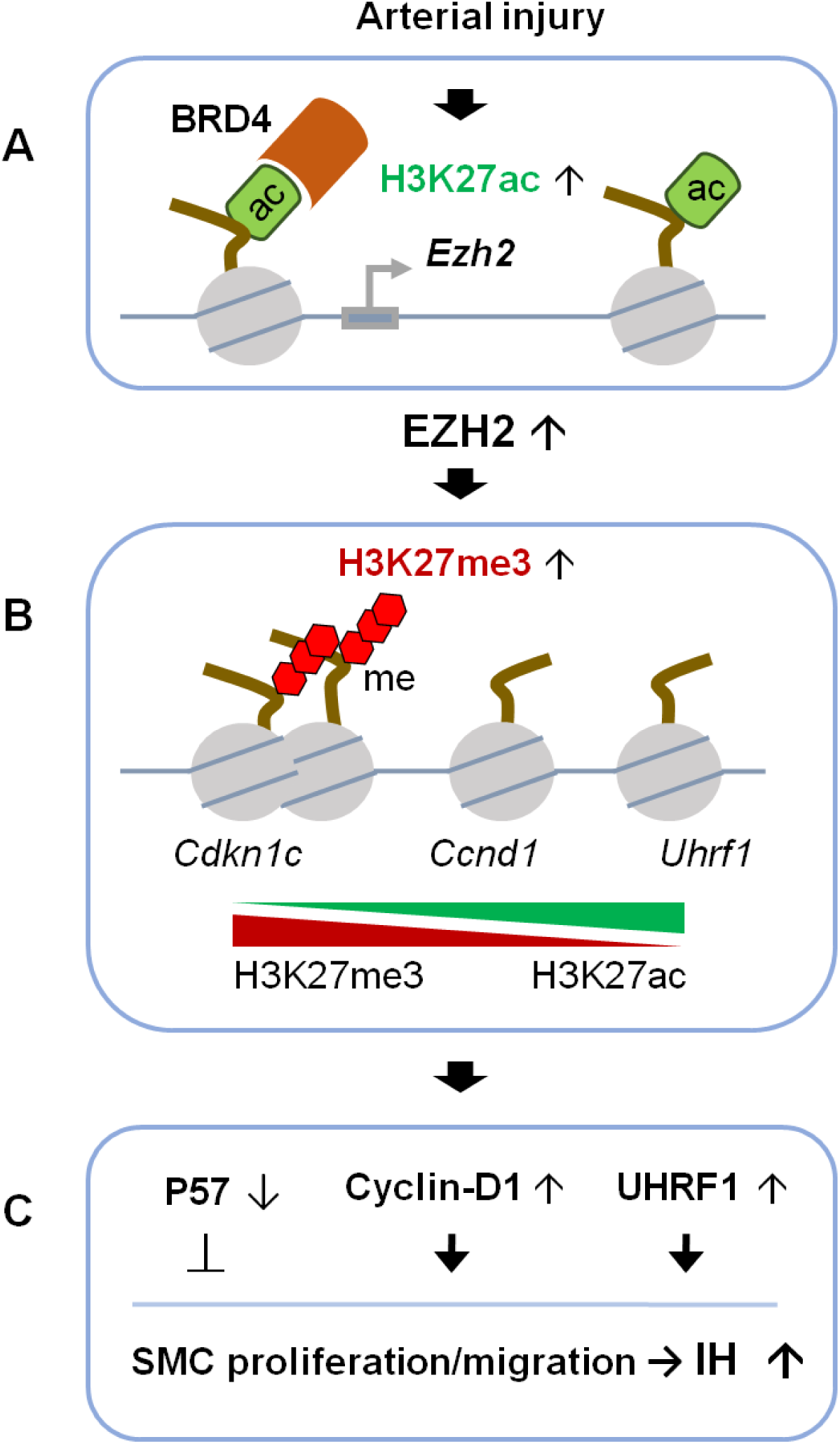
Schematic depiction of BRD4 regulation of EZH2 expression and function. Angioplasty injury in rat carotid arteries induces BRD4/H3K27ac enrichment at *Ezh2*, upregulation of EZH2 protein, H3K27me3 accumulation at *Cdkn1c* and attenuation at *Ccnd1*, and exacerbation of IH.

## Conclusions

The current study integrated information from in vivo epigenomic mapping, conditional knockout and gene transfer in rodent models of IH, as well as in vitro mechanistic elaboration. The results together with our previous reports^3, 6, 10^ depicted a coherent story with new insights. In response to angioplasty injury, BRD4, a histone acetylation reader and transcription co-activator^26^, clustered with H3K27ac and enhancers at *Ezh2*. Up-regulated EZH2 may effectuate H3K27me3 redistribution from activators to inhibitors of SMC proliferation. The observed functional consequences manifested as enhanced pro-IH SMC behaviors and exacerbated IH. Thus, the herein released new knowledge may help anti-IH translational design, now that clinical tests targeting BRD4 and EZH2 in other conditions have heralded progress^14^.

## Methods

### Animals

All animal studies conform to the *Guide for the Care and Use of Laboratory Animals* (National Institutes of Health) and protocols approved by the Institutional Animal Care and Use Committee at The Ohio State University (Columbus, Ohio).

### Balloon angioplasty in rat carotid arteries

To induce IH, the Fogarty balloon catheter for clinical thrombectomy (2F, Edwards Scientific) was applied in male Sprague-Dawley rats (300 to 350 g) to injure the left common carotid artery, as we previously described^6^. Briefly, an incision was made in the neck of anesthetized animal. Through an opening on the left external carotid artery, the balloon was inserted and advanced ∼1.5 cm into the common carotid artery, inflated (at 1.5 atm), withdrawn to the bifurcation, and then deflated before next insertion. This procedure was repeated 3 times. Blood flow was resumed in the common and internal carotid arteries (after ligating the external artery). The animal was maintained in general anesthesia with inhalation of 2-2.5 % of isoflurane. Analgesics including carprofen, bupivacaine, and buprenorphine were injected to the animal recovering from anesthesia. Animals were euthanized in a chamber slowly filled with CO_2_.

### Artery tissue ChIP sequencing and data processing

To preserve the artery “real-time” epigenetic information, balloon-injured and uninjured (contralateral) common carotid arteries were snap frozen in liquid N_2_ immediately after dissected and severed out. Artery collection was performed 7 days after balloon angioplasty. Artery tissues from 40 rats were pooled for ChIPseq analysis at Active Motif per company standard procedures. Briefly, chromatin was isolated after adding lysis buffer, followed by disruption with a Dounce homogenizer. Genomic DNA was sheared to an average length of 300–500 bp by sonicating the lysates, and the segments of interest were immunoprecipitated using an antibody (4μg) against BRD4, H3K27a, H3K27me3, or H3K4me1. The protein/DNA complexes eluted from beads were treated with RNase and proteinase K, crosslink was reversed, and the ChIP DNA was then purified for use in the preparation of Illumina sequencing libraries. Standard steps included end-polishing, dA-addition, adaptor ligation, and PCR amplification. The DNA libraries were quantified and sequenced on Illumina’s NextSeq 500, as previously described^26^. Sequence reads were aligned to the reference genome Rn5, peak locations were identified using Macs2 algorithm^33^ and annotated based on UCSC RefSeq. Differential peak locations were called using SICER^34^. In-house shell and R scripts (https://www.r-project.org) were used for data integration. To summarize and cluster genome-wide TSS coverage as heat maps, deepTools (PMID: 24799436) compute matrix and plotheatmap functions were utilized. IGV (http://www.broadinstitute.org/igv/) was used for visualization. Annotation files were downloaded from UCSC.

### Conditional knockout of BRD4 and mouse femoral artery wire injury

The *Brd4*^*fl/fl*^ mouse line^35^ with loxP sites flanking *Brd4* exon 3 were kindly provided by Dr. Keiko Ozato from National Institute of Child Health and Human (NICHD). The smooth muscle lineage-specific, tamoxifen-inducible Cre strain (*Myh11-CreER*^*T2*^) was purchased from The Jackson Laboratory. These two strains were crossed, and the offsprings carrying *Brd4*^*fl/fl*^ and/or *Myh11-CreER*^*T2*^ were selected through RT-PCR genotyping as previously described^35^. Genotyping PCR primers are provided in Table S3. Mice were fed with tamoxifen-citrate chow (TD.130860) for 10 days, and then with normal diet for another 7 days prior to femoral artery wire injury to induce IH.

Mouse femoral artery wire injury was performed as described in detail in our publication dedicated to this model^36^. Briefly, a midline incision was made in the ventral left thigh to dissect the common femoral artery. The distal and proximal ends of the femoral artery were temporally looped. An arteriotomy was made on the deep femoral artery muscular branch, through which a 0.015″ guide wire (REF#C-SF-15-15, Cook Medical, Bloomington, IN) was inserted and kept stationary for 1 minute. After removal of the wire, the muscular branch was ligated and blood flow was resumed. At 28 days after injury, femoral arteries were collected following perfusion fixation (with PBS first and then 4% paraformaldehyde) at a physiological pressure of 100 mmHg. The animal was kept anesthetized with inhalation of 5% of isoflurane throughout the terminal procedure. Animals were euthanized in a chamber slowly filled with CO_2_.

### Lentiviral vector construction for EZH1 or EZH2 silencing and overexpression

To construct a lentiviral vector for the expression of EZH1- or EZH2-specific shRNAs, the pLKO.1-puro empty vector was purchased from Addgene (Watertown, MA). A scrambled shRNA control and shRNAs specific for the mouse EZH1 and EZH2 genes were designed by RNAi Central (http://cancan.cshl.edu/RNAi_central/step2.cgi). The corresponding shRNA-expressing lentivectors were constructed by using the pLKO.1-puro vector as a template.

Lentiviruses were packaged in Lenti-X 293T cells (cat#632180, Clontech, Mountain View, CA) using a three-plasmid expression system (pLKO.1-shRNAs-puro, psPAX2 and pMD2.G) as described in our recent reports^6, 10^, and used in combination (5:3:2) Efficient sequences (based on siRNAs as final products) are listed in Table S1.

### Intraluminal infusion of lentivirus and perivascular inhibitor drug delivery

To express a transgene or shRNA, lentivirus was infused into the balloon-injured artery wall as we recently described in detail. Briefly, immediately after angioplasty, a cannula was inserted through the external carotid artery arteriotomy, advanced past the bifurcation, and ligated to generate a sealed intraluminal space in the common carotid artery. A syringe containing lentivirus was connected to the cannula. The virus (total 150 ul, >1×10^9^ IFU/ml) was slowly injected, incubated for 25min in the lumen. The lumen was then flushed repeatedly with saline containing 20U/ml heparin and blood flow resumed. Heparin was also administered perioperatively to prevent thrombosis.

For pharmacological local treatment of injured rat carotid arteries, a thermosensitive hydrogel (AK12, Akina Inc., IN) was used for perivascular administration of the EZH1/2 inhibitor UNC1999, following our published method. Briefly, immediately after angioplasty, UNC1999 (10 mg/rat) or an equal amount of DMSO (vehicle control) dispersed in 400 μl AK12 gel was applied around the balloon-injured artery. The surgery was then finished as described above for the angioplasty model.

### Morphometric analysis of IH and restenosis

Paraffin sections (5 μm thick) were cut using a microtome (Leica) at equally spaced intervals and then stained (van Gieson or hematoxylin and eosin) for morphometric analysis, as described in our previous reports. Morphometric parameters as follows were measured on the sections and calculated by using ImageJ software: area inside external elastic lamina (EEL area), area inside internal elastic lamina (IEL area), lumen area, intima area (= IEL area - lumen area), and media area (= EEL area – IEL area). Intimal hyperplasia (IH) was quantified as a ratio of intima area versus media area (I/M). Measurements were performed by an independent researcher blinded to the experimental conditions using 3 to 6 sections from each of rat. The data from all sections were pooled to generate the mean for each animal. The means from all the animals in each treatment group were then averaged, and the SEM was calculated.

### Immunoblotting

Cells or rat carotid artery homogenates (pulverized in liquid nitrogen) were lysed in radio-immunoprecipitation assay (RIPA) buffer containing protease inhibitors (50 mM Tris, 150 mM NaCl, 1% Nonidet P-40, 0.1% sodium dodecyl sulfate, and 10 μg/ml aprotinin). Approximately 15-30 μg of proteins from each sample were separated via sodium dodecyl sulfate-polyacrylamide gel electrophoresis on a 10% gel. The proteins were then transferred to a polyvinylidene difluoride membrane and detected by immunoblotting. The antibody sources and dilution ratios are listed in Table S2. Specific protein bands on the blots were illuminated by applying enhanced chemiluminescence reagents (Thermo Fisher Scientific; Catalog no. 32106) and then recorded with an Azur LAS-4000 Mini Imager (GE Healthcare Bio-Sciences, Piscataway, New Jersey). Band intensity was quantified by using ImageJ software.

### Assays for proliferation and migration

Proliferation was determined by using the CellTiter-Glo Luminescent Cell Viability kit (Promega, Madison, Wisconsin) following manufacturer’s instructions. Wildtype or lentiviral infected MOVAS (a mouse vascular smooth muscle line) cells were seeded in 96-well plates at a density of 2,000 cells per well with a final volume of 200 μl DMEM (10% FBS). Cells were then starved with 0.5% FBS overnight and then stimulated with PDGF-BB (20 ng/ml). At 72 h of PDGF-BB treatment, plates were decanted, refilled with 50 μl CellTiter-Glo reagent/50 μl phosphate-buffered saline per well, and incubated at room temperature for 10 min before reading in a FlexStation 3 Benchtop Multi-Mode Microplate Reader (Molecular Devices, San Jose, California) (250-ms integration).

To determine cell migration, scratch (wound healing) assay was performed as described in our previous report. Briefly, wildtype or lentiviral-infected MOVAS cells were cultured to a 90% confluency in 6-well plates and then starved overnight. A sterile pipette tip was used to generate an ∼1 mm cell-free gap. Dislodged cells were washed away with PBS. Plates were then refilled with fresh medium containing 20 ng/ml of PDGF-BB and incubated for 24 h. Calcein AM was then added (2 μM) to illuminate the cells. After a 15-min incubation, cells were washed 3 times with PBS, and images were then taken. Cell migration was quantified by ImageJ software (National Institutes of Health, Bethesda, Maryland) based on the change in the width of the cell-free gap before and after PDGF-BB stimulation.

### Quantitative real-time polymerase chain reaction (qPCR)

Assays were performed following our published methods. Briefly, total ribonucleic acid was isolated from cultured cells or rat carotid arteries (pulverized in liquid nitrogen) by using a Trizol reagent (Thermo Fisher Scientific) following the manufacturer’s protocol. Potential contaminating genomic deoxyribonucleic acid (DNA) was removed by using gDNA Eliminator columns provided in the kit. Total ribonucleic acid of 1 μg was used for the first-strand complementary DNA synthesis (Thermo Fisher Scientific). Quantitative real-time polymerase chain reaction was performed by using Quant Studio 3 (Thermo Fisher Scientific). Each complementary DNA template was amplified in triplicate PerfeCTa SYBR Green SuperMix (Quantabio, Beverly, Massachusetts). Primers are listed in Table S3.

### Statistical Analysis

Data are presented as mean ± standard error of the mean (SEM). For statistical analysis, one-way ANOVA followed by post-hoc Tukey’s test was applied for multi-group comparison and unpaired Student t-test (and also Mann–Whitney non-parametric test) was used for 2-group comparison, as specified in each figure legend. Statistical significance was set at P< 0.05. For ChIPseq data, ***s***tatistical analyses were performed using SAS/STAT software, version 9.2 (SAS Institute, Inc., Cary, NC).

## Acknowledgement

This work was supported by NIH grants R01 HL133665 (to L.-W. G.), R01HL-143469 and R01HL-129785 (to K.C.K., L.-W. G.), and an AHA pre-doctoral award 17PRE33670865 (to M.X.Z.). We thank Dr. Keiko Ozato (Section on Molecular Genetics of Immunity, NICHD, NIH) for kindly providing the *Brd4*^*fl/fl*^ mouse strain.

## Figure legends

**Table S1.**
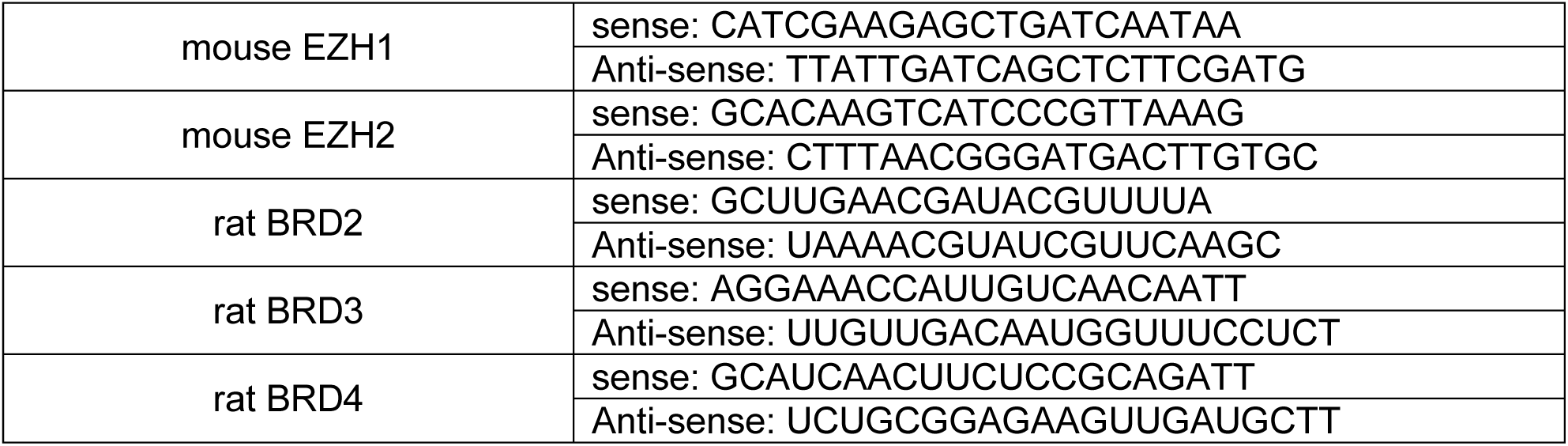
siRNA or shRNA sequences for mouse and rat genes.

**Table S2.** Antibodies.

| Antigen | Manufacturer | Catalog Number | Dilution Ratio | Application |
| --- | --- | --- | --- | --- |
| H3K27me3 | Cell Signaling Technology | 9733 | 1:1000 | Western Blot |
| H3K27me3 | Cell Signaling Technology | 9733 | 1:100 | Immunofluorescence |
| EZH1 | Proteintech | 20852-1-AP | 1:500 | Western Blot |
| EZH2 | Cell Signaling Technology | 5246 | 1:1000 | Western Blot |
| BRD4 | Abcam | Ab128874 | 1:1000 | Western Blot |
| CyclinD1 | Cell Signaling Technology | 55506 | 1:1000 | Western Blot |
| P57 | Cell Signaling Technology | 2557 | 1:1000 | Western Blot |
| β-actin | Abcam | Ab8226 | 1:5000 | Western Blot |

**Table S3.**
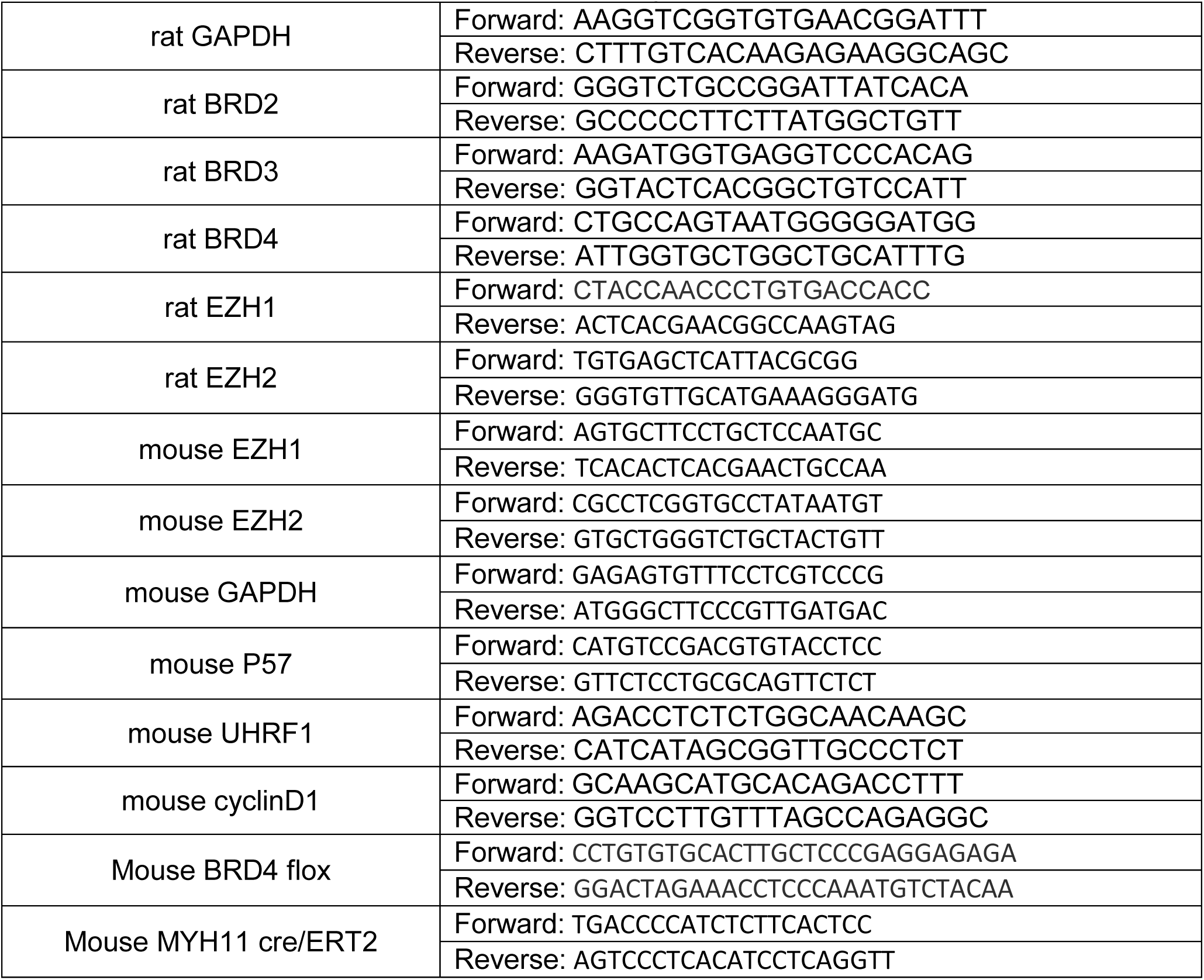
Primer sequences for mouse and rat genes.

